# Single- and duplex TaqMan-quantitative PCR for determining the copy numbers of integrated selection markers during site-specific mutagenesis in *Toxoplasma gondii* by CRISPR-Cas9

**DOI:** 10.1101/2022.06.23.497358

**Authors:** Kai Pascal Alexander Haenggeli, Andrew Hemphill, Norbert Müller, Bernd Schimanski, Philipp Olias, Joachim Müller, Ghalia Boubaker

## Abstract

Herein, we developed a single and a duplex TaqMan quantitative qPCR for absolute quantification of copy numbers of integrated dihydrofolate reductase-thymidylate synthase (*mdhfr-ts*) drug selectable marker for pyrimethamine resistance in *Toxoplasma gondii* knockouts (KOs). The single TaqMan qPCR amplifies a 174 bp DNA fragment of the inserted *mdhfr-ts* and of the wild-type (WT) *dhfr-ts* (*wt-dhfr-ts*) which is present as single copy gene in *Toxoplasma* and encodes a sensitive enzyme to pyrimethamine. Thus, the copy number of the *dhfr-ts* fragment in a given DNA quantity from KO parasites with a single site-specific integration should be twice the number of *dhfr-ts* copies recorded in the same DNA quantity from WT parasites. The duplex TaqMan qPCR allows simultaneous amplification of the 174 bp *dhfr-ts* fragment and the *T. gondii 529-bp* repeat element. Accordingly, for a WT DNA sample, the determined number of tachyzoites given by *dhfr-ts* amplification is equal to the number of tachyzoites determined by amplification of the *Toxoplasma 529-bp*, resulting thus in a ratio of 1. However, for a KO clone having a single site-specific integration of *mdhfr-ts*, the calculated ratio is 2. We then applied both approaches to test *T. gondii* RH mutants in which the *major surface antigen* (SAG1) was disrupted through insertion of *mdhfr-ts* using CRISPR-Cas9. Results from both assays were in correlation showing a high accuracy in detecting KOs with multiple integrated *mdhfr-ts*. Southern blot analyses using BsaBI and DraIII confirmed qPCRs results. Both TaqMan qPCRs are needed for reliable diagnostic of *T. gondii* KOs following CRISPR-Cas9-mediated mutagenesis, particularly with respect to off-target effects resulting from multiple insertions of *mdhfr-ts*. The principle of the duplex TaqMan qPCR is applicable for other selectable markers in *Toxoplasma*. TaqMan qPCR tools may contribute to more frequent use of WT *Toxoplasma* strains during functional genomics.

## Introduction

*Toxoplasma gondii* is an apicomplexan parasite that causes diseases in farm animals with an enormous global economic impact and a high zoonotic potential [1]. In immunocompetent hosts, infection does not have serious consequences, and proliferative tachyzoites differentiate into tissue cyst-forming bradyzoites, which can persist over many years to lifelong without causing any clinical symptoms. However, *T. gondii* is an opportunistic pathogen, and primary infection in pregnant animals and also women can lead to vertical transmission, and result in fetal malformations and/or abortion. In patients undergoing immunosuppression, either by disease or through medical treatment, reactivation of bradyzoites from tissue cysts and re-differentiation into tachyzoites frequently causes serious pathology. Current drugs for toxoplasmosis treatment typically include antifolates using a combination of pyrimethamine–sulfadiazine or trimethoprim– sulfamethoxazole, and pyrimethamine can also be combined with clindamycin, azithromycin, or atovaquone. These treatments are unspecific, frequently result in adverse effects, and clinical failures have been reported [2,3]. To date, more than 110 years after the first discovery of *T. gondii* [4], there is still a need for identifying drug targets and vaccine candidates, which could be exploited for the development of better preventive or therapeutic options for the management of toxoplasmosis (Blader *et al*., 2015; Sidik *et al*., 2016). In this context, functional genomics plays a major role, and gene knockout (KO) in protozoan parasites is the most commonly applied approach [7]. *Toxoplasma* is highly amenable to genetic manipulation and has thus emerged as one of the major apicomplexan model parasites [8].

Gene KO and gene replacement strategies rely on double crossover homologous recombination (HR) using type I or II *T. gondii* KU80 mutants (*Δku80*s) as parental strain [9]. The *Δku80* parasites are deficient in the non-homologous end-joining (NHEJ) pathway required for repairing DNA double-strand breaks (DSBs) [10,11]. Genetic manipulation of *T. gondii* WT strains is hindered by the presence of a predominant NHEJ as main DSB repair mechanism [12], which results in enhanced random integration of exogenous genes. Despite the fact that *Δku80* background increases the efficiency of targeted mutagenesis in *T. gondii* by HR, defective NHEJ might render parasites prone to accumulate chromosomal aberrations [13] causing genomic instability [14], in particular since *T. gondii* tachyzoites are usually maintained *in vitro* through excessive cycles of proliferation and DNA replication.

For positive selection of *T. gondii* mutant or transgenic strains that have successfully integrated an exogenous DNA coding for a modified dihydrofolate reductase-thymidylate synthase (mDHFR-TS), pyrimethamine (Pyr) is the drug of choice [15–19], since mDHFR-TS confers resistance to Pyr [20]. In the genome of WT *T. gondii*, a single-copy gene coding for DHFR-TS (WT-DHFR-TS) is expressed, but the enzyme is sensitive to Pyr [21]. The mDHFR-TS differs from WT-DHFR-TS by three amino acid substitutions; two are in exon 1 (Ser **TC**T → Arg **CG**T and Thr A**C**C → Asn A**A**C) and one in exon 3 (Phe **T**TT → Ser **T**CC) [20–22].

Based on the original method of CRISPR-Cas9 that was successfully implemented for genome editing in *T. gondii* in 2014 [23,24], many alternative protocols have been developed [9] rendering genetic manipulation of WT strains feasible. This advance was possible because in CRISPR-Cas9 formation of a DSB at a specified genomic site is ensured by the 20-nucleotide guide RNA (gRNA) that binds and guides the Cas9 endonuclease to the defined location [25]. Then, the CRISPR-Cas9 mediated DNA break can be repaired through NHEJ or homology-directed repair (HDR) pathways [9]. Options for delivering CRISPR-Cas9 components into the cell as one- or two-vector or cloning-free approaches are now available [26].

Although CRISPR-Cas9 has significantly improved the efficiency of targeted mutagenesis and/or site-specific insertion of selectable markers in *Toxoplasma* WT strains, the *Δku80* parasites have remained the first choice for functional genetic studies [9,27–29]. For *Δku80* strains, CRISPR-Cas9 has allowed to considerably reduce the length of homologous flanking DNA to 40 bps [23]. This has rendered the task of template DNA preparation more simple, since these short homology flanking regions of 40 bps can be incorporated into primers designed for the amplification of the selectable marker [26].

A crucial step during CRISPR-Cas9 is the verification of the KO and the validation of gene edits, which must be carried out prior to further functional investigations. Following the selection of mutant clones by drug treatments, PCR and/or Sanger sequencing are used to verify the DNA sequence of the targeted locus [23,26]. Subsequently, Western blotting and/ or immunostaining are applied to confirm the loss of gene expression [23]. Nonetheless, off-target effects (OTEs) of CRISPR-Cas9 are often not considered. OTEs resulting from non-specific cleavage by a uncomplexed Cas9 are of low probability, since endonuclease activity of Cas9 is dependent on the interaction with the gRNA [30] as revealed by crystallographic studies [31–35]. However, a gRNA-independent endonuclease activity by Cas9 in the presence of Manganese ions was reported [36].

Overall, the gRNA and the protospacer adjacent motif (PAM) next to the targeted genomic sequence play a crucial role in determining the specificity of gene targeting by CRISPR-Cas9 [37]. For example, the *Streptococcus pyogenes* Cas9 (*Sp*Cas9) binds optimally to a consensus NGG canonical PAM [38–40], but it can also interact, albeit with less affinity, with other non-canonical PAMs [41] mostly NAG and NGA [42,43]. Furthermore, Cas9 can unspecifically cleave a DNA sequence with up to seven mismatch base pairs in the PAM proximal region of the gRNA sequence known as “seed sequence” [44,45]. In addition, in the mammalian genome, DNA or RNA bulges, caused by small insertions or deletions, were identified as potential off-target sites [46]. The incidence of off-target mutations by CRISPR-Cas9 widely varies between cell types and species [37], particularly in cells with defective DSB repair pathways [47]. Substantial efforts have been made to develop *in silico* systems for optimal gRNA design. However, prediction and scoring by the algorithms employed are mostly based on DNA-binding rather than cleavage, and even more significant factors such as PAMs, DNA/RNA bulges and experimental conditions are excluded [48].

Whole genome sequencing (WGS) is the only unbiased and direct approach allowing a comprehensive analysis of OTEs including single-nucleotide polymorphisms, indels and other structural differences. However, this approach is costly and time consuming, thus cannot be applied as a first-line testing strategy [37]. Moreover, when the designed strategy to achieve gene KO by CRISPR-Cas9 consists in disrupting the targeted sequence followed by insertion of a selectable marker, it is important to check KO cells for unintended additional integration events. For that, Southern blotting (SB) can be applied, which allows to determine the copy number of inserted exogenous DNA. However, SB requires a relatively large amount of DNA, special equipment, and is relatively time-consuming when many clones have to be analyzed. In addition, the accuracy of SB depends largely on the use of appropriate restriction enzymes.

An alternative strategy to determine single or multiple transgene integration events caused by CRISPR-Cas9 is real-time PCR-based quantification (RT-qPCR), which allows a more high-throughput determination of transgene copy numbers and respective integration patterns (single or multiple insertions) [49–52].

In this study, we aimed at improving the selection protocol for *T. gondii* KO transfectants generated by CRISPR-Cas9, with regard to the identification of OTEs resulting from multiple insertion of selectable marker by developing two TaqMan qPCR-based approaches

## Materials and methods

### Parasite and cell culture

Tachyzoites of *T. gondii* type I RH strain were maintained *in vitro* in human foreskin fibroblasts (HFF) as previously described [53].

### CRISPR-Cas9 compounds and *mdhfr-ts* selection cassette

The DNA sequence coding for *T. gondii* RH SAG1 was retrieved from GenBank under the accession number GQ253075.1 and used for the design of the 23-nt gRNA (Table 1).

**Table 1:**
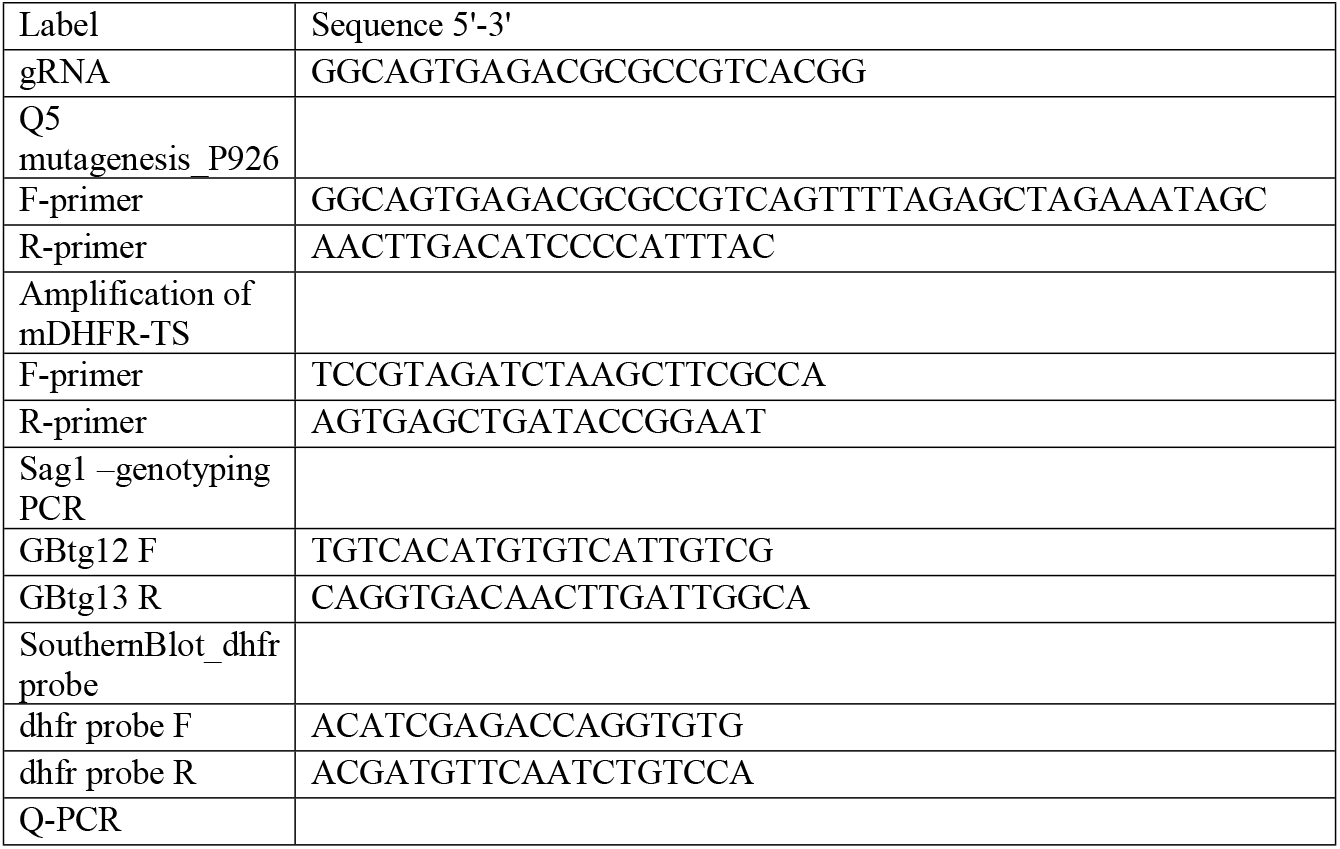
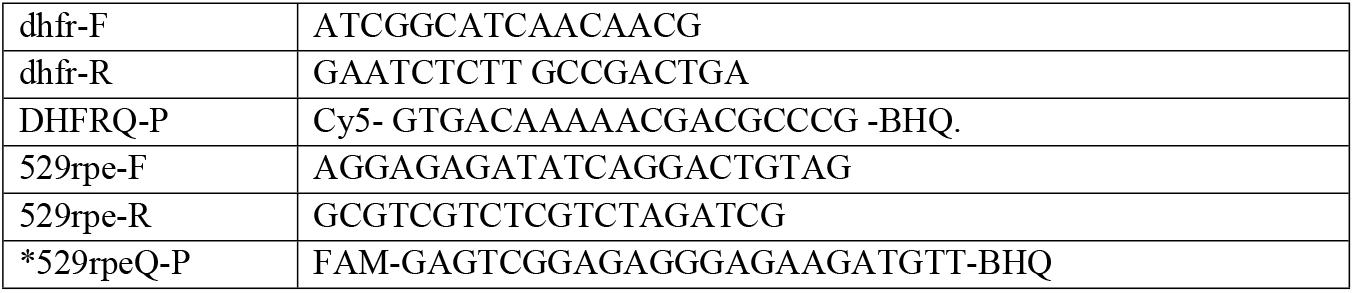
Sequence of primers and probes used in this study (*) TaqMan probes designed in this study.

The plasmid P926 encodes a GFP tagged Cas9 endonuclease and a pre-existing gRNA. The expression of Cas9 is under the control of the bacterial T7 promotor while transcription of the gRNA is driven by the *T. gondii* U6 promotor [23]. The pre-existing gRNA was replaced a by the newly designed DNA using the Q5 Site-Directed mutagenesis kit (New England Biolabs, M0491S), respective primers are listed in Table 1. The resulting plasmid was then amplified in NEB 5-alpha competent *Escherichia coli* (*E. coli*), purified using ZymoPURE Plasmid Miniprep Kit (Zymo Research) and sequenced. The plasmid P972 was used for amplification of the selectable marker *mdhfr-ts*.

### Transfection and selection

The transfection procedure was adapted from Sidik et al. [23]. Briefly, the electroporation reaction was prepared in a final volume of 300 µL cytomix buffer containing 7.5 μg P926, 1.5 μg of *mdhfr-ts*, 0.112 × 10^7^ *T. gondii* RH WT tachyzoites, 2μM adenosine triphosphate (ATP) and 5μM L-glutathione in 4 mm gap cuvettes (Axonlab, Baden, Switzerland). Cells were than electroporated with a pulse generator (ECM830, BTX Harvard Apparatus, Holliston, MA) by applying the following protocol: 1700 V, 176*μ*s of pulse length, two pulses with 100 ms interval. Transfected tachyzoites were transferred immediately into T25 flasks with confluent HFFs, which were placed in a humidified incubator at 37°C / 5%-CO_2_. After 24 h, the cultures were subjected to drug selection by the addition of 3µM Pyr to the culture medium. Clones were isolated by limiting dilution (0.5 tachyzoites/150 µL medium) and allowed to grow in 96 well plates for 10 days.

### PCR and Sanger sequencing

Genomic DNA from thirty-three clones and WT tachyzoites extracted using NucleoSpin DNA RapidLyse kit (Macherey-Nagel) according to the manufacturer’s instructions. We further genotyped the SAG1 locus of the thirty-three clones by PCR. Amplicons of the WT SAG1 locus were □ 216 bp, however, for KO clones with one insertion of the complete MDHFR-TS sequence, the expected amplicon length was 3379 bp (∼3400 bp). The diagnostic PCR was performed in 50 µL final volume containing 0.2 mM dNTPs, 0.5 µM of each forward (GBtg12) and reverse primers (GBTg13), Q5 high-fidelity DNA polymerase (1 unit) and Q5 high GC enhancer (1x), and 80 ng of template DNA. The GBtg12 F/ GBtg13 R primer sequences are shown in Table 1. Conditions were as follows: initial denaturation at 98°C for 3 min; 25 cycles of denaturation at 98°C for 30 sec, annealing at 58°C for 30 sec, and elongation at 72°C for 2 min. The final cycle was followed by extension at 72°C for 2 min. PCR products were purified using Zymo DNA Clean and Concentrator kit (Zymo Research), 20 ng of purified PCR products were submitted to Sanger sequencing.

### Immunofluorescence assay (IFA)

Immunofluorescence microscopy was done as described previously [54,55]. Briefly, freshly egressed tachyzoites were isolated from infected HFF cultures, fixed in suspension in PBS/3% paraformaldehyde, and were allowed to attach to poly-L-lysine-coated coverslips for 20 min at room temperature. To permeabilize cells, coverslips were incubated with pre-cooled methanol/acetone (1:1) solution for 20 min at −20 °C. Then samples were rehydrated and incubated overnight at 4°C in PBS/3 % bovine serum albumin (BSA) solution to block unspecific binding sites. SAG1 expression was assessed by using α-SAG1 monoclonal antibody (1:1000) and anti-mouse fluorescein-isothiocyanate (FITC) (1:300). For double stainings, SAG1 labelled parasites were further incubated in polyclonal rabbit anti-Inner Membrane Complex 1 (IMC1) antibody (1:500), and a secondary anti-rabbit tetramethyl-rhodamine-isothiocyanate (TRITC) (1:300). Finally, cover slips were mounted onto glass slides using Vectashield mounting medium containing 4, 6-diamidino-2-phenylindole (DAPI).

### SDS-PAGE and Western Blotting

Pellets corresponding to equal numbers of WT or *Δsag1* tachyzoites were prepared and dissolved in Laemmli SDS sample buffer, which contains β-mercaptoethanol. Cell lysates were then separated by SDS-PAGE. Two SDS-PAGEs were made simultaneously; after electrophoresis, one gel was stained with Coomassie and proteins on the other gel were transferred to nitrocellulose filters. The blot was saturated with blocking solution (5% skimmed milk powder and 0.3% Tween 20 in PBS) for 2 hours at room temperature and then incubated with *T. gondii* α-SAG1 monoclonal antibody (1:500) overnight at 4°C. After washing, nitrocellulose membrane was incubated with an alkaline-phosphatase conjugated anti-mouse IgG antibody (1:1000). Lastly, reactive bands were visualized by immersion of the blot in 5-bromo-4-chloro-3-indolyl phosphate (BCIP)/nitro blue tetrazolium (NBT) detection solution.

### Single TaqMan-qPCR

To determine the copy numbers of the inserted *mdhfr-ts* selectable marker in the genome of KO clones, we designed a single TaqMan-qPCR taking advantage of the fact that WT *T. gondii* tachyzoites have a single copy of *dhfr-ts* in their genome (*wt-dhfr-ts*). Specific *dhfr* forward and reverse primers (Table 1) were designed to yield a 174 bp fragment of the MDHFR-TS or WT-DHFR-TS gene. The TaqMan probe DHFRQ-P (Table 1) contained the Cyanine 5 (Cy5) reporter dye at the 5′ end and Black Hole Quencher (BHQ) fluorescent quencher at the 3′ end.

Freshly egressed tachyzoites from infected cultures were filtered through a 3-μM pore-sized polycarbonate membrane, counted and 10^6^ tachyzoites were used for DNA extraction by NucleoSpin DNA RapidLyse Kit according to the instructions provided by the manufacturer. From each tested WT or KO clone, 3 ng DNA were used as template. DNA quantifications were performed by QuantiFluor double-stranded DNA (dsDNA) system (Promega, Madison, WI, USA). PCR amplification was performed in a total reaction mixture of 10 μL containing 1x SensiFast master mix (Bioline, Meridian Bioscience), 0.5 μM of reverse and forward primers, 0.1 μM of DHFRQ-P probe, 0.3 mM dUTP, and one unit of heat-labile Uracil DNA Glycosylase (UDG) [56]. A Bio-Rad CFX 96 QPCR instrument (Biorad) was used with the following thermal profile: (1) initial incubation of 10 min at 42 °C, followed by (2) denaturation step of 5min at 95°C and (3) 50 cycles of two-step amplification (10 s at 95 °C and 20 s at 62 °C). Samples were tested in triplicates and a negative control with double-distilled water was included for each experiment. For quantification, two standard curves were made: one was based on the use of a 10-fold serial dilution of the plasmid P972 ranging from 1.29 × 10^9^ to 1.29 copies/ 3 μl, and the other one was based on a 10-fold serial dilution of DNA from WT *T. gondii* RH, with tachyzoite numbers ranging from 7.5 × 10^5^ to 75 per 3 μL [57].

### Duplex TaqMan-qPCR

In this assay, the number of tachyzoites corresponding to 3ng DNA and the copy number of the DHFR-TS DNA fragment were assessed simultaneously. Quantification of tachyzoites was achieved by preparation of a *T. gondii* standard curve using 10-fold serial dilutions with parasite concentrations ranging from 7.5 × 10^5^ to 75 and amplification of a 162-bp region of the *T. gondii* 529-bp repeat element [58]. Amplifications were carried-out in total volume of 10 µL containing 1x SensiFast master mix (Bioline, Meridian Bioscience), 0.5 μM of each primer set (dhfr-F/R and 529rpe-F/R), 0.1 μM of each probe (DHFRQ-P and 529rpeQ-P), 0.3 mM dUTP, and one unit of heat-labile uracil DNA glycosylase (UDG). From each sample, three ng of DNA were used in the reaction mix. All reactions were run in triplicates and amplifications were carried-out under the same thermal profile used for the single TaqMan-qPCR. The cycle threshold values (CT) were plotted as mean of triplicates against the standard curve values to determine the number of tachyzoites. Parasite concentrations were determined after the calculation of the linear regression equation (y = ax + b), where y = CT; a = curve slope (slope); x = parasite number; and b = where the curve intersects y-axis (y intercept).

### Southern Blot

Two Southern hybridizations were carried out on seven *Δsag1* clones and the WT strain that were tested by qPCRs. One µg of each genomic DNA-sample was digested with the restriction enzymes BsaBI or DraIII for 6 h at 60°C or 37°C, respectively. Reaction mixtures were then separated by 0.8% agarose gel electrophoresis containing ethidium bromide. Gels were subjected to depurination (15 min in 0.25 M HCl), denaturation (30 min in 1M NaCl/0.5 M NaOH) and neutralization (1 hour in 1M Tris-HCl, pH 7.5/ 3 M NaCl). Separated DNA fragments were then transferred onto Hybond membrane (Amersham) by capillary transfer and subsequently stably fixed by UV crosslinking for 10 seconds. For blocking non-specific binding sites, membranes were pre-incubated in hybridization buffer (0.5 M Na_2_HPO_4_, 60 mM H_3_PO_4_, 7% SDS, 1% BSA, 0.9 mM EDTA) for 2 hours at 65°C.

The DHFR probe was generated from the plasmid P972 by PCR with DHFR forward and reverse primers listed in Table 1, gel-purified and radioactively labelled with α-P32-dCTP using the Amersham Megaprime DNA Labeling System. The labeled probe was heat-denatured at 95°C for 3 min and added directly to the pre-hybridized membranes. After overnight incubation at 65°C the membranes were washed 15 minutes each in 1xSSC, 0.1% SDS and 0.5xSSC, 0.1% SDS and eventually exposed to Phosphoimager screens for 20 hours.

## Results

### Generation of *T. gondii* RH *Δsag1* clones by CRISPR/Cas9

After transfection and 10 days *in vitro* culture under Pyr treatment, thirty-three clones, together with WT parasites were genotyped by PCR. As shown in Figure 1 (Fig. 1), the WT locus produced the expected PCR product of □ 216 bp. Five clones, namely *T. gondii* RH *Δsag1* C18, 23, 30, 31 and 33 exhibited a PCR product in the expected size of more than 3kb, indicating integration of the selection marker. In other clones such as in *T. gondii* RH *Δsag1* C6 and C7, PCR amplified a product of ≤ 1000 bp.

**Fig 1.**
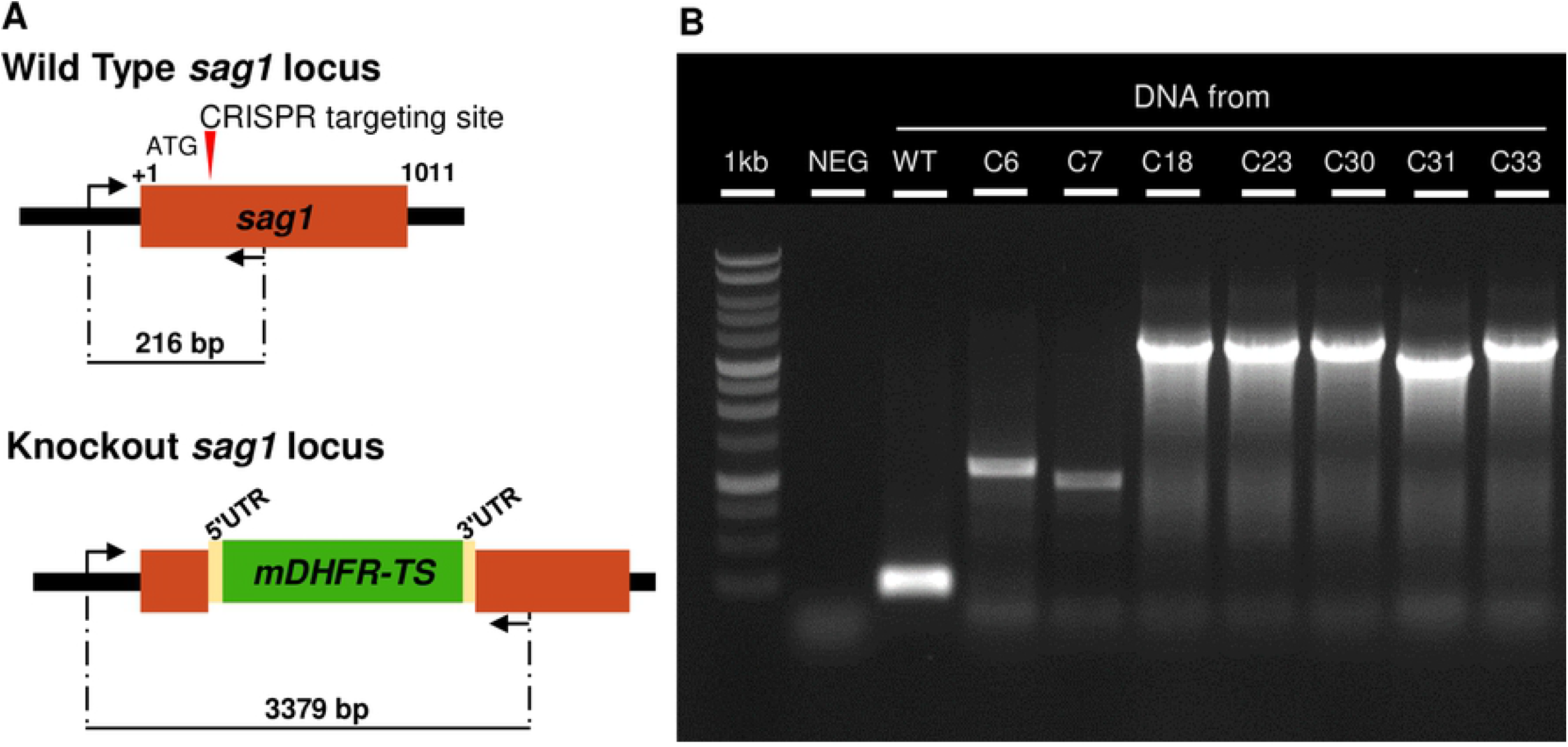
SAG1 gene disruption in *T. gondii* RH by CRISPR-Cas9 technology. (**A**) Schematic representation of the strategy used to disrupt *sag1* by inserting the pyrimethamine-resistance gene MDHFR-TS. (**B**) Diagnostic PCR revealing integration of a complete *mdhfr-ts* sequence into *sag1* in four clones (C18, C23, C30 and C33) compared with the parental strain RH. The KO clone C31 showed a smaller band, clones C6 and 7 exhibited a band ≤ 1000 bp. The WT locus produced the expected PCR product (□ 216 bp).

Direct Sanger sequencing of the obtained PCR products revealed that in *T. gondii* RH *Δsag1* C18, 23, 30 and 33, *sag1* was disrupted by insertion of complete *mdhfr-ts* sequence, while clone C31 had incorporated a truncated *mdhfr-ts* into *sag1*. For clone 6 and 7, the DSB in the SAG1 gene generated by CRISPR-Cas9 was repaired through NHEJ by insertion of short DNA sequence, while the actual selection marker *mdhfr-ts* was most likely integrated elsewhere in the genome. Western blot analysis as well as IFA confirmed the absence of TgSAG1 expression in tachyzoites of *T. gondii RH Δsag1* C6, 7, 18, 23, 30, 31 and 33 (supplementary Fig. 1).

### Single TaqMan-qPCR

As shown in Figure 2 (Fig. 2), this single TaqMan-qPCR aimed to determine whether random integration in *T. gondii RH Δsag1* C18, 23, 30, 31 and 33 occurred elsewhere in the genome beside the detected site-specific integration of *mdhfr-ts* in *sag1*. The principle is based on the fact that the copy number of the *dhfr-ts* fragment in a given DNA quantity of KO parasites with a single site-specific integration should be twice the number of *dhfr-ts* copies recorded in the same DNA quantity from WT parasites (Fig. 2).

**Fig 2.**
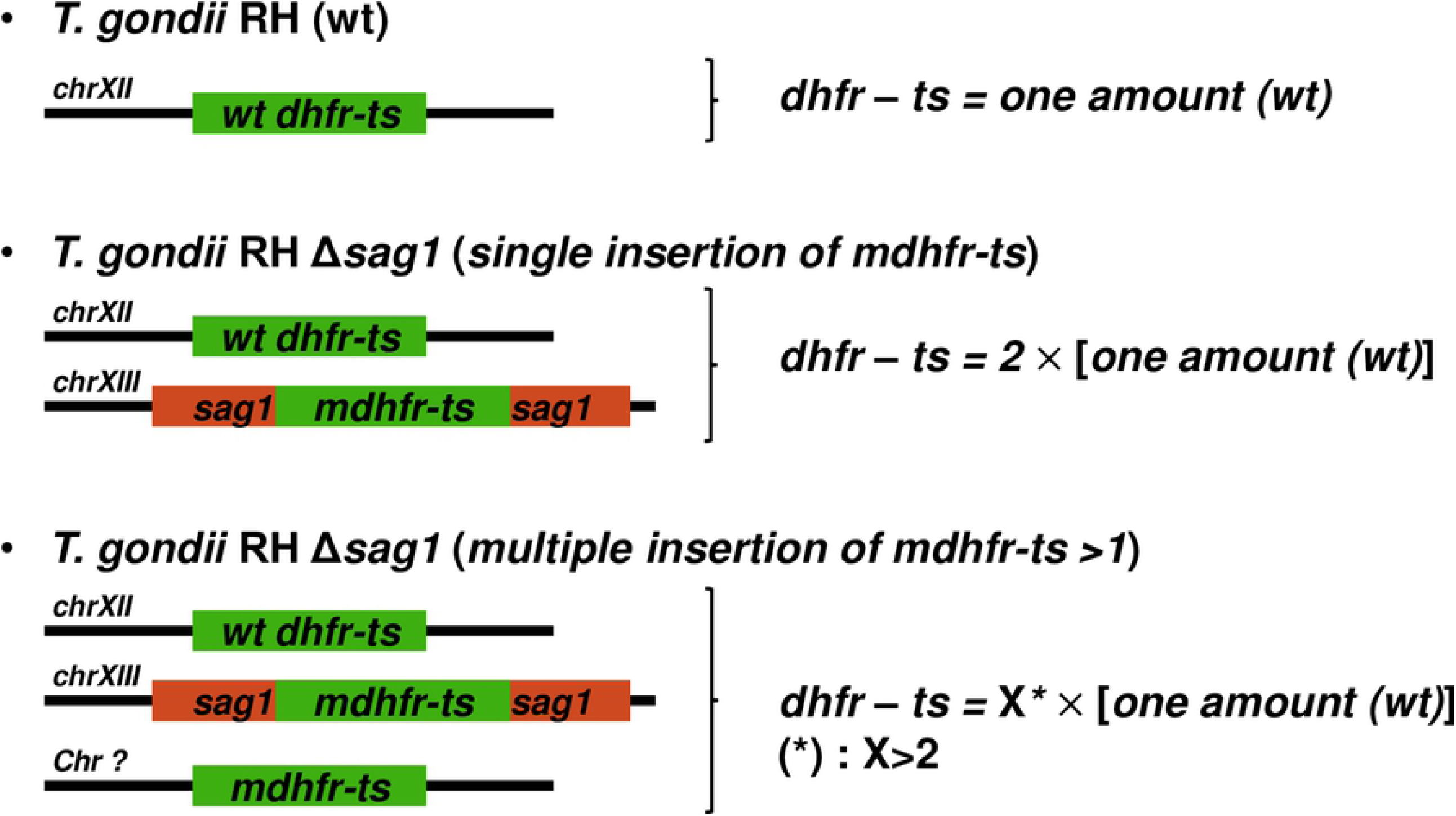
Principle and potential outcomes of the single TaqMan-qPCR.

As shown in Figure 3 A and B, comparable linear calibrator curves were obtained using serial 10-fold dilutions of *mdhfr-ts* plasmid or *T. gondii* genomic DNA (range 7.5 × 10^5^ to 75 genome equivalents), indicating thus similar amplification efficiency of *dhfr-ts* from both sources (Fig. 3 A and B).

**Fig 3.**
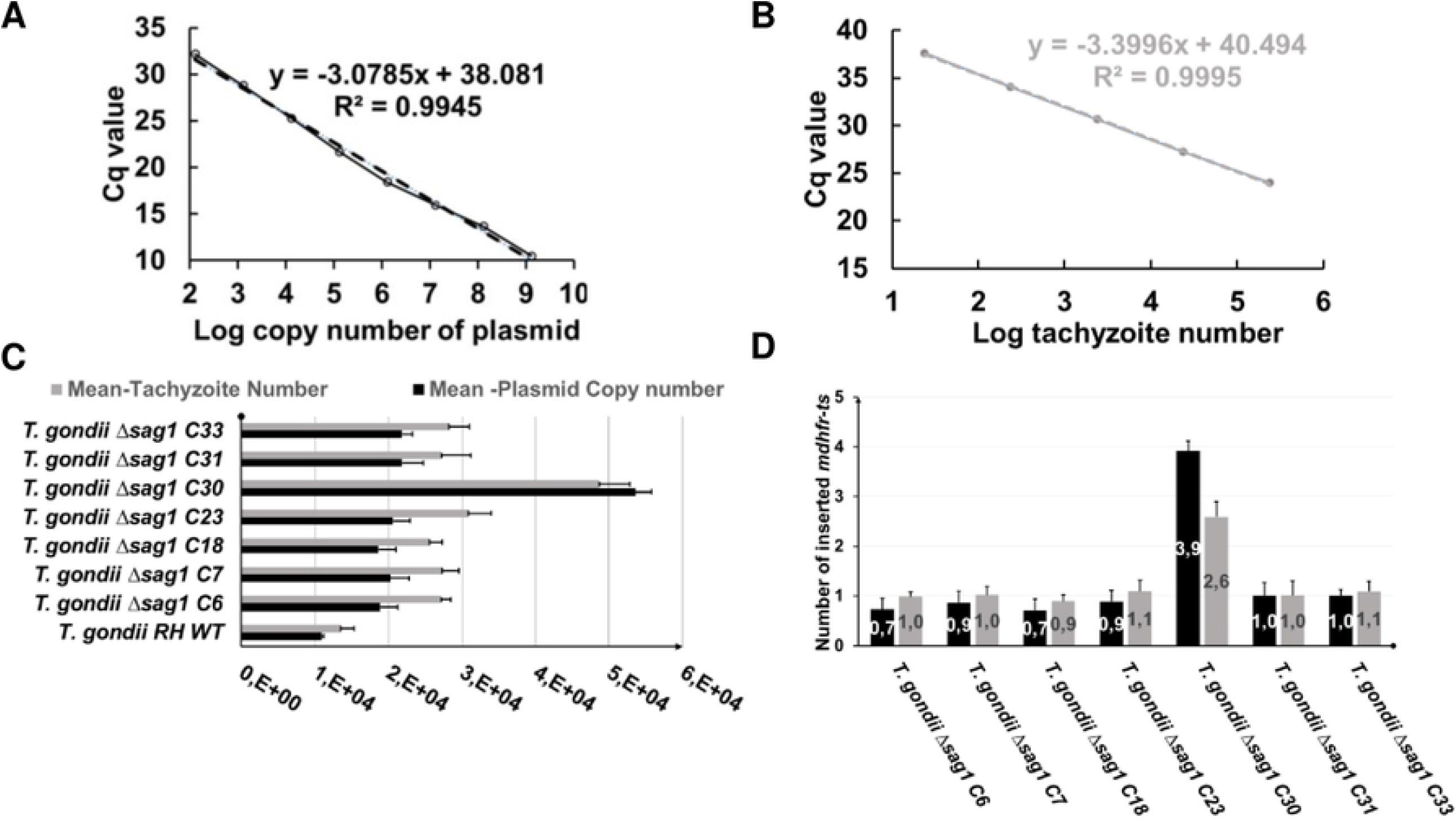
Single TaqMan-qPCR for determining the copy number of integrated *mdhfr-ts* selectable marker. Standard curves were made through a triplicate test of 10-fold serial dilutions of (**A**) P972 or (**B**) *T. gondii* RH DNA. (**C**) For each WT or KO clone, the number of existing *dhfr-ts* in the genome was determined according to the plasmid based standard curve (black bars) and the *T. gondii* RH DNA-based calibrator (grey bars). Since in the *T. gondii* genome the *wtdhfr-ts* is a single copy gene, the following equation was used: one WT tachyzoite = one-copy *dhfr-ts*, for the calculation based on *T. gondii* RH DNA based calibrator curve (grey bars). Error bars indicate standard deviation of triplicates for each sample. In (**D**), the number of inserted *mdhfr-ts* in each KO clone is defined by subtracting the *dhfr-ts* copy number found in the WT from the *dhfr-ts* copy number in the KO (black bars) or by subtracting the tachyzoite numbers determined for the WT from tachyzoite numbers corresponding the KO clone (grey bars). The optimal result of 1 indicates a single integration event of the *mdhfr-ts* into *sag1*.

As shown in Figure 3 C, for clones *T. gondii* RH *Δsag1* C6, 7, 18, 23, 31 and 33, the determined number of *dhfr-ts* copies in the three ng of DNA was almost the double of that number calculated for the WT parasites, independently of the used standard curve. The calculated number of inserted *mdhfr-ts* selectable marker was almost equal to 1 for the following clones: *T. gondii* RH *Δsag1* C6, 7, 18, 23, 31 and 33, as shown in Figure 3 D (Fig. 3 D).

### Duplex TaqMan-qPCR

In this assay, quantitative amplification of the *dhfr-ts* and of the *T. gondii* 529-bp repeat element were combined into one reaction (Fig. 4).

**Fig 4.**
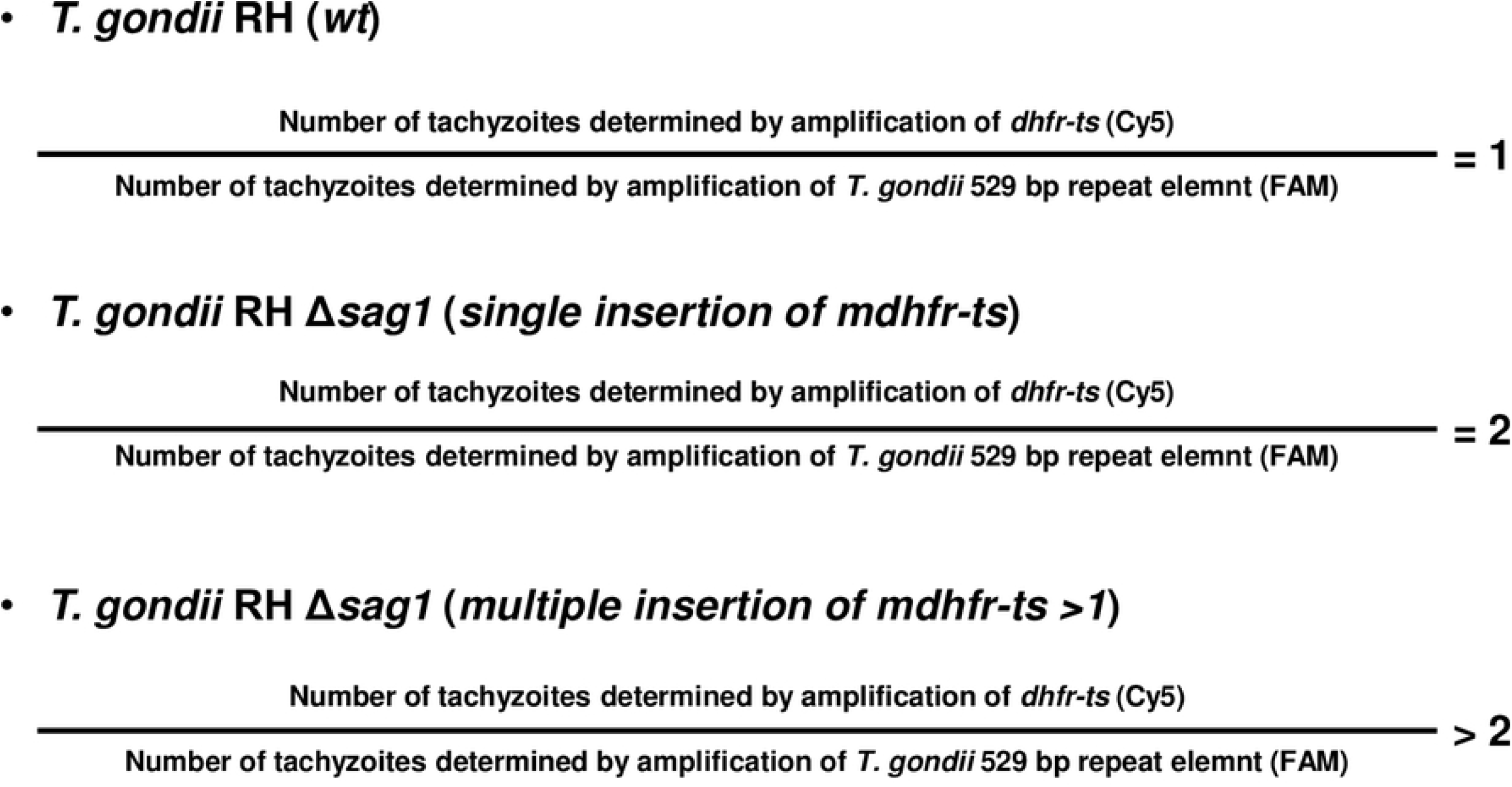
Principle and potential outcome of the duplex TaqMan-qPCR.

The standard curve was made from a 10-fold serial dilution of *T. gondii* RH DNA, with parasite concentrations ranging from 7.5 × 10^5^ to 75 ^(^Fig. 5 A). The designed two primer pairs in the duplex TaqMan-qPCR enabled similar amplification efficiencies (R2 = 0.99 %) for their respective targets (Fig. 5 A).

**Fig 5.**
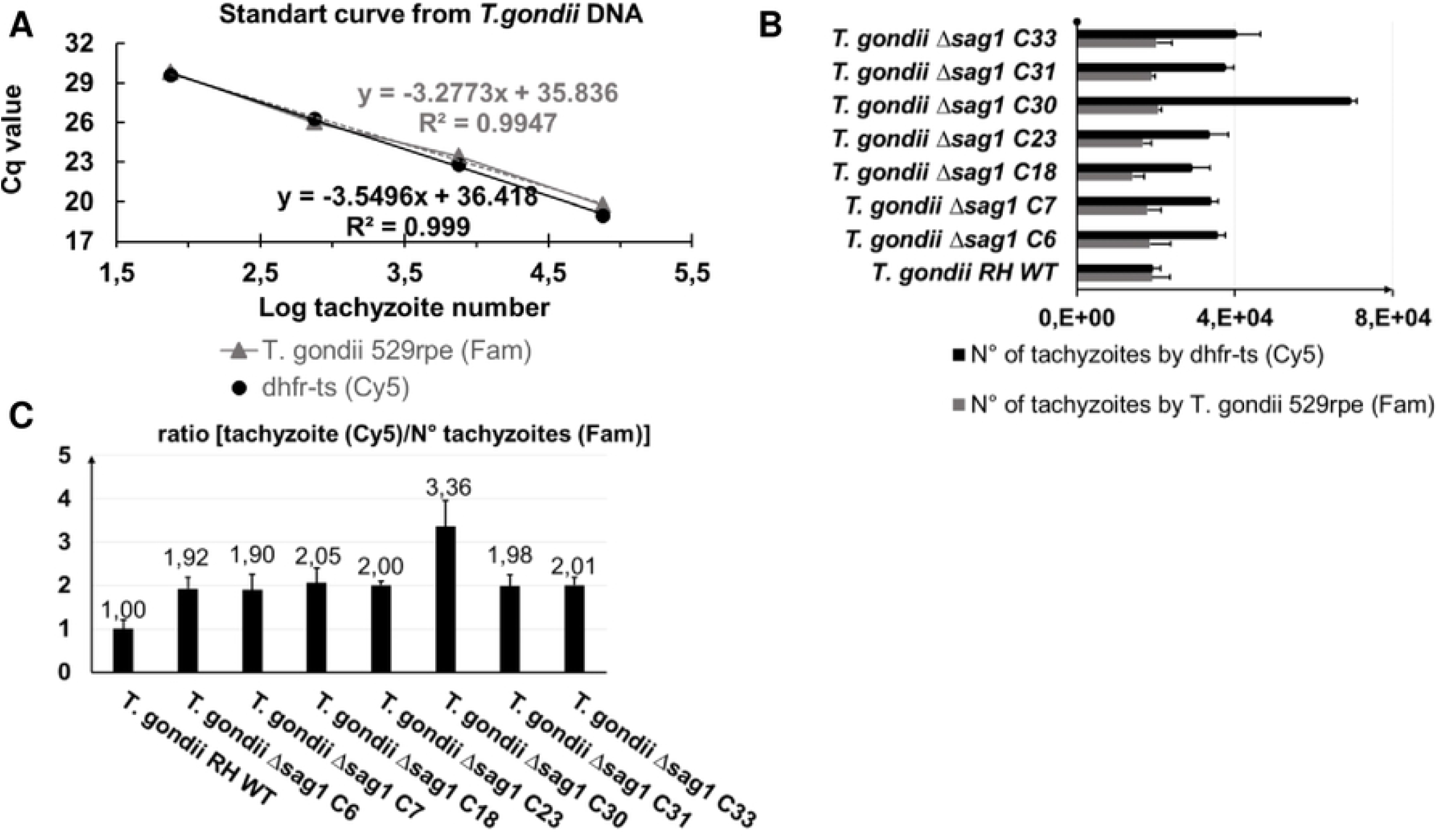
Duplex TaqMan-qPCR for determining copy numbers of integrated mdhfr-ts selectable marker. (**A**) Standard curve was made by using a 10-fold serial dilution of *T. gondii* RH DNA, with tachyzoites numbers ranging from 75 to 7.5 × 10^5^ parasites. (**B**) For each WT or KO clone, the numbers of tachyzoites in the 3 ng DNA was determined according to amplification of *dhfr-ts* (black bars) and to the *T. gondii* 529 bp repeat element (grey bars). In (**C**), the number of inserted *mdhfr-ts* is given by the ratio of the number of tachyzoites as determined by *dhfr-ts* amplification and the number of tachyzoites determined by using the *T. gondii* 529 bp repeat element. A ratio equal to 2 indicates a single integration event of the *mdhfr-ts* in *sag1*. Error bars indicate standard deviations of triplicates for each sample

For the wild type DNA sample, the determined number of tachyzoites given by *dhfr-ts* amplification is equal to the number of tachyzoites determined by amplification of the *Toxoplasma* 529-bp (Fig. 5 B), resulting thus in a ratio of 1 (Fig. 5 C). For a *T. gondii* RH *Δsag1* clone having a single insertion of the *mdhfr-ts* within *sag1*, the calculated ratio is estimated to be 2, as it is the case for clone C6, 7, 18, 23, 31 and 33 (Fig. 5 C). For *T. gondii* RH *Δsag1* C30, the number of tachyzoites given by *dhfr-ts* quantification was more than three times higher than the number of tachyzoites obtained by amplification of the *Toxoplasma* 529-bp repeat element (ratio > 3), which is indicative for multiple insertions of the *mdhfr-ts* fragment into the genome (Fig. 5 C).

### Southern blot analysis

To validate the results from both single and duplex TaqMan-qPCRs concerning the numbers of integrated *mdhfr-ts* fragments into the genome, Southern blot analysis of genomic DNA digested with BsaBI and DraIII was carried out (Fig. 6). In the case of BsaBI digestion, (Fig. 6 A) the labeled probe recognized a 14.177-kb fragment in the *wtdhfr-ts* gene and a 4.999-kb fragment in the *mdhfr-ts* selectable marker integrated into *sag1*, such that the integrated fragment is easily identified in *sag1* KO parasites (Fig. 6 A). For genomic DNA digested with DraIII, the *wtdhfr-ts* is present in all clones at 5.164 kb, and the integrated *mdhfr-ts* fragment within *sag1* is found at 4.351 kb (Fig. 6 B).

**Fig 6.**
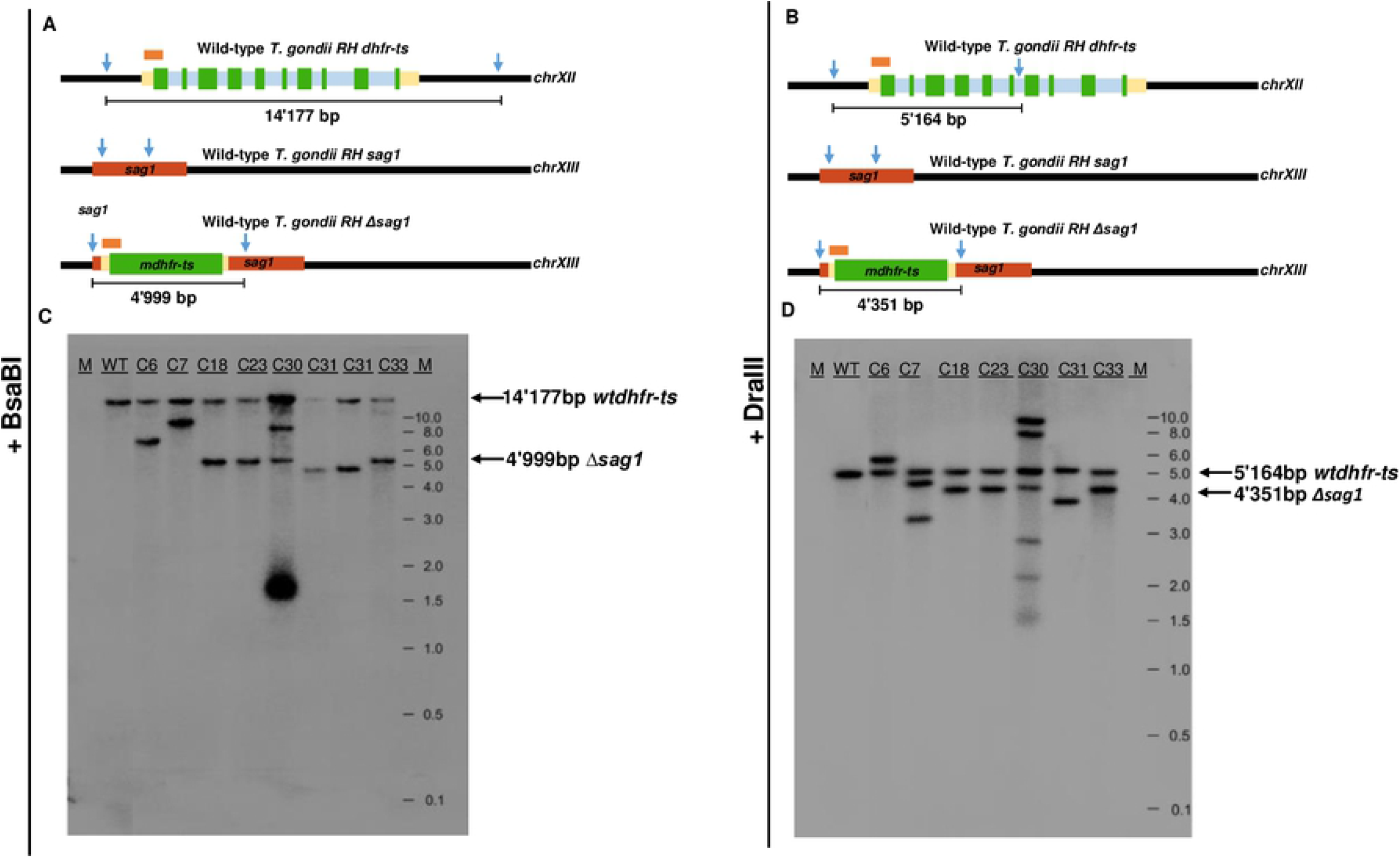
Southern blot analysis for determining the number of mdhfr-ts integration events into the *T. gondii* RH genome. (**A**) and (**B**) Schematic drawing of hybridization probe and restriction sites of BsaBI and DraIII in the WT *T. gondii* RH *dhfr-ts* gene and in the WT and mutant *sag1* locus. (**C**) Southern blot of genomic DNA digested with BsaBI and (**D**) with DraIII. M indicates the size of the fragments separated by gel electrophoresis.

SB showed that WT *T. gondii* RH, as well as all seven KO clones, exhibited a single band corresponding to the *wtdhfr-ts* gene, migrating at 14.17 kb in BsaBI-digested DNA and at 4.99 kb in DraIII-digested DNA. Clones *T. gondii* RH *Δsag1* C18, 23 31 and 33 exhibited a two-band pattern after digestion with BsaBI or DraIII, confirming thus a single integration event of the *mdhfr-ts* selection marker in the genome (Fig. 6 C and D). The band hybridizing with the probe in DNA of clone *T. gondii* RH *Δsag1* C31 was at a lower position than the one observed for C18, 23 and 33. Thus, in agreement with the sequencing analysis, the inserted selectable marker within *sag1* in clone C31 is a truncated version of *mdhfr-ts*. For *T. gondii RH Δsag1* C6, SB also revealed an integration of only one *mdhfr-ts* copy into the genome, but at another position than the *sag1* gene. This was also the case for *T. gondii RH Δsag1* C7, with the exception that after genomic DNA digestion with DraIII, three bands were found to be hybridizing with the probe. Concerning the clone *T. gondii* RH *Δsag1* C30, additional hybridizations were detected after digestion with BsaBI (two bands) or DraIII (four bands) besides the expected *wtdhfr-ts* and *mdhfr-ts* bands, indicating random and multiple integrations of *mdhfr-ts* into the *T. gondii* RH genome.

## Discussion

In this study, we have established a single- and duplex TaqMan-qPCR assay for determination of copy numbers of integrated *mdhfr-ts* selectable marker to evaluate of *T. gondii* RH KO parasites generated by CRISPR-Cas9 as exemplified by using the major tachyzoite surface antigen TgSAG1 as KO target gene. *T. gondii* RH *Δsag1* clones lacking the expression of TgSAG1 generated through CRISPR-Cas9 mediated KO were selected by treatment with Pyr, and the lack of TgSAG1 expression was ascertained by IFA and WB. Considering the risks of OTEs and thus the random integration of gene fragments into the genome, the *sag1* locus in different clones was amplified by PCR and respective fragments were sequenced to assess integration of the *mdhfr-ts* selection marker. A single- and duplex Taq Man qPCR for determination of the copy numbers of *mdhfr-ts* in *T. gondii* RH *Δsag1* tachyzoites was developed, and was validated by SB.

Using CRISPR-Cas9 technology, the efficiency of *sag1* disruption through insertion of the *mdhfr-ts* selectable marker without homology arms was about 15 % (5/33). Thus CRISPR-Cas9 increased gene editing efficiency in WT *Toxoplasma* compared to a frequency a 2×10^−5^ obtained by non-homologous recombination [59]. The efficiency of 15% obtained herein can be considered satisfactory since WT *Toxoplasma* strains are significantly more relevant for studying gene function than most commonly used NHEJ-deficient *Δku80* strains. So far, frequency, severity, and the types of DNA sequence changes that might occur in association with the lack of NHEJ in *Δku80* parasites remains largely unknown. In apicomplexan parasites particularly *Theileria parva, Cryptosporidium spp*. and *Plasmodium spp*., loss of the classical NHEJ (C-NHEJ) pathway over genome evolution is suggested to be associated with reduced genome size (8 - 23 megabytes), this in comparison to the *T. gondii* genome (87 megabytes) that encodes the three main components of the C-NHEJ namely Ku70, Ku80 and DNA ligase IV [60]. In eukaryotic cells, impaired DNA-DSB repair pathways contributes to significant stress-induced effects and causes genomic instability [13,14,61]. Moreover, the use of *Δku80* strains for functional genomics does not prevent hazardous insertion of exogenous donor DNA. For example, cases of random integration into the genome were reported during reverse genetics in malaria parasites [62] naturally lacking key NHEJ compounds [63,64].

For CRISPR-Cas9, OTEs resulting from non-targeted DNA mutations (base substitutions, deletions and insertions) are of low probability; in hematopoietic stem- and progenitor cells, rate of insertion–deletion mutations did not differ between Cas9-treated and non-Cas9-treated cells [65]. These results were reported from two independent experiments targeting two different genes located in different chromosomes [65]. Thus, for reliable transgenesis and genome editing in *Toxoplasma* using selectable markers, selection protocols of engineered cells must include a step for determining whether an unintended integration of exogenous DNA has occurred.

Despite Southern blot analysis is ranked second after the WGS as the most unambiguous method for estimation of copy number in transgenic unicellular protozoan parasites; it has also significant disadvantages; particularly it is unsuitable for automation since the choice of restriction enzymes and probes are experiment-specific. Furthermore, digestion with restriction enzymes may result in DNA fragments larger than 15 kb, which are inefficiently blotted, leading thus to an underestimated copy number.

In contrast to SB, qPCR can be used to scan the entire genome for the presence of a selectable marker independently of the genomic location, and this can be done at higher throughput and in a wide dynamic range, which in turn allows simultaneous testing large numbers of samples in a short time frame. Consequently, qPCR was successfully implemented as an alternative to SB for characterization of transgene copy number and integration site in many different transgenic plant and animal cells [66,67]. In this study, the strong evidence in line with this recommendation is *T. gondii* RH *Δsag1* C30, which would have been taken for a correct mutant without further evaluation by single- and duplex TaqMan-qPCR, which detected multiple insertions. In addition, results for *T. gondii* RH *Δsag1* C6 and C7 clearly demonstrate that both TaqMan-qPCRs can provide an absolute quantification of the inserted selection marker, independently of its location in the genome. This was in line with PCR-Sequencing and SB findings, demonstrating a single copy integration of *mdhfr-ts* elsewhere in the genome for both *T. gondii* RH *Δsag1* C6 and C7. Concerning KO C7, the appearance of two bands in SB upon digestion with DraIII, but not with the BsaBI restriction enzyme, together with the results of the qPCRs, strongly suggest that the insertion of the single copy *mdhfr-ts* in an unknown genomic location has generated a new cutting site for DraIII.

Regarding the quantification of inserted *mdhfr-ts* copies in the examined clones, results obtained with the single TaqMan-qPCR were in correlation with those resulting from duplex TaqMan-qPCR. Thus, both single and duplex TaqMan-qPCR protocols can be applied as described herein each time *mdhfr-ts* is chosen as a selection marker in *Toxoplasma* gene KO experiments. So far, *mdhfr-ts* has been the most commonly used selection marker for transgenic *T. gondii* and *P. falciparum* [20,68].

The duplex TaqMan-qPCR presented here can also be employed in case other selection markers are chosen. In such a case, primers and probes specific to the amplification of the *Toxo 529-bp repeat element* can be used as reported here, however new primers and a TaqMan probe specific to the exogenous DNA needs to be designed. Subsequently, two important aspects need to be considered: (i) both primer sets must result in similar amplification efficiency; (ii) the standard curves must be made using *Toxoplasma* parasites as reference that have only one copy of the designed selection marker. Positive selection strategies based on drug resistance are limited in *T. gondii*; thus besides the *mdhfr* resistance gene [20], choices are almost restricted to *E. coli* chloramphenicol acetyl transferase (*cat*) [69,70] or *Streptoalloteichus* ble (*ble*) [71] genes, which confer resistance to chloramphenicol or phleomycin, respectively. In order to ensure the maximum accuracy of single and duplex TaqMan-qPCR results, standardized protocols for cell-culture, tachyzoite purification, DNA extraction and quantification should be applied to all tested mutants/clones.

In conclusion, we have developed and validated sensitive, rapid and reliable single and duplex TaqMan qPCR methods for measuring *mdhfr-ts* copy numbers during CRISPR-Cas9 mediated gene editing in *Toxoplasma*. A significant advantage of these quantitative assays, particularly the duplex TaqMan qPCR, is that they can be easily applied for any selection cassette other than *mdhfr-ts*. Therefore, we believe that both qPCR techniques could become methods of choice for characterizing transgenic *T. gondii* cell-lines in term of integration pattern of the used exogenous DNA. Furthermore, by providing such a versatile molecular tools for quantitative detection of the integrated selection cassette, WT *T. gondii* stains can now be more frequently used instead of *ku80* KO strains.

## Acknowledgements

Anti-SAG1 and anti-IMC1 antibodies used in this study were a kind gift from Prof. Dominique Soldati-Favre, University of Geneva.

---

**Supplementary Figure 1**. Loss of *sag1* expression in *T. gondii* RH SAG1 knockouts by (**A**) Western blot analysis and (**B**) immunofluorescence.

## References

1. Matta SK, Rinkenberger N, Dunay IR, Sibley LD. Toxoplasma gondii infection and its implications within the central nervous system. Nat Rev Microbiol. 2021;19: 467–480. doi:10.1038/s41579-021-00518-7

2. Alday PH, Doggett JS. Drugs in development for toxoplasmosis: advances, challenges, and current status. Drug Des Devel Ther. 2017;11: 273–293. doi:10.2147/DDDT.S60973

3. Dunay IR, Gajurel K, Dhakal R, Liesenfeld O, Montoya JG. Treatment of Toxoplasmosis: Historical Perspective, Animal Models, and Current Clinical Practice. Clin Microbiol Rev. 2018;31: e00057–17. doi:10.1128/CMR.00057-17

4. Nicolle C, Manceaux LH. On a leishman body infection (or related organisms) of the gondi. 1908. Int J Parasitol. 2009;39: 863–864. doi:10.1016/j.ijpara.2009.02.001

5. Blader I, Coleman B, Chen C-T, Gubbels M-J. The lytic cycle of Toxoplasma gondii: 15 years later. Annu Rev Microbiol. 2015;69: 463–485. doi:10.1146/annurev-micro-091014-104100

6. Sidik SM, Huet D, Ganesan SM, Huynh M-H, Wang T, Nasamu AS, et al. A Genome-wide CRISPR Screen in Toxoplasma Identifies Essential Apicomplexan Genes. Cell. 2016;166: 1423-1435.e12. doi:10.1016/j.cell.2016.08.019

7. Ren B, Gupta N. Taming Parasites by Tailoring Them. Front Cell Infect Microbiol. 2017;7: 292. doi:10.3389/fcimb.2017.00292

8. Oberstaller J, Otto TD, Rayner JC, Adams JH. Essential Genes of the Parasitic Apicomplexa. Trends Parasitol. 2021;37: 304–316. doi:10.1016/j.pt.2020.11.007

9. Di Cristina M, Carruthers VB. New and emerging uses of CRISPR/Cas9 to genetically manipulate apicomplexan parasites. Parasitology. 2018;145: 1119–1126. doi:10.1017/S003118201800001X

10. Fox BA, Ristuccia JG, Gigley JP, Bzik DJ. Efficient gene replacements in Toxoplasma gondii strains deficient for nonhomologous end joining. Eukaryot Cell. 2009;8: 520–529. doi:10.1128/EC.00357-08

11. Fox BA, Falla A, Rommereim LM, Tomita T, Gigley JP, Mercier C, et al. Type II Toxoplasma gondii KU80 Knockout Strains Enable Functional Analysis of Genes Required for Cyst Development and Latent Infection. Eukaryot Cell. 2011 [cited 5 Jan 2022]. doi:10.1128/EC.00297-10

12. Fenoy IM, Bogado SS, Contreras SM, Gottifredi V, Angel SO. The Knowns Unknowns: Exploring the Homologous Recombination Repair Pathway in Toxoplasma gondii. Front Microbiol. 2016;7: 627. doi:10.3389/fmicb.2016.00627

13. Difilippantonio MJ, Zhu J, Chen HT, Meffre E, Nussenzweig MC, Max EE, et al. DNA repair protein Ku80 suppresses chromosomal aberrations and malignant transformation. Nature. 2000;404: 510–514. doi:10.1038/35006670

14. Nussenzweig A, Sokol K, Burgman P, Li L, Li GC. Hypersensitivity of Ku80-deficient cell lines and mice to DNA damage: The effects of ionizing radiation on growth, survival, and development. Proc Natl Acad Sci. 1997;94: 13588–13593. doi:10.1073/pnas.94.25.13588

15. Wang L, Tang D, Yang C, Yang J, Fang R. Toxoplasma gondiiADSL Knockout Provides Excellent Immune Protection against a Variety of Strains. Vaccines. 2020;8: 16. doi:10.3390/vaccines8010016

16. Ma Z, Yan K, Jiang R, Guan J, Yang L, Huang Y, et al. A Novel wx2 Gene of Toxoplasma gondii Inhibits the Parasitic Invasion and Proliferation in vitro and Attenuates Virulence in vivo via Immune Response Modulation. Front Microbiol. 2020;11: 399. doi:10.3389/fmicb.2020.00399

17. Zheng J, Cheng Z, Jia H, Zheng Y. Characterization of aspartyl aminopeptidase from Toxoplasma gondii. Sci Rep. 2016;6: 34448. doi:10.1038/srep34448

18. Yang W-B, Wang J-L, Gui Q, Zou Y, Chen K, Liu Q, et al. Immunization With a Live-Attenuated RH:ΔNPT1 Strain of Toxoplasma gondii Induces Strong Protective Immunity Against Toxoplasmosis in Mice. Front Microbiol. 2019;10: 1875. doi:10.3389/fmicb.2019.01875

19. Li M, Liu J, Wu Y, Wu Y, Sun X, Fu Y, et al. Requirement of Toxoplasma gondii metacaspases for IMC1 maturation, endodyogeny and virulence in mice. Parasit Vectors. 2021;14: 400. doi:10.1186/s13071-021-04878-0

20. Donald RG, Roos DS. Stable molecular transformation of Toxoplasma gondii: a selectable dihydrofolate reductase-thymidylate synthase marker based on drug-resistance mutations in malaria. Proc Natl Acad Sci U S A. 1993;90: 11703–11707. doi:10.1073/pnas.90.24.11703

21. Roos DS. Primary structure of the dihydrofolate reductase-thymidylate synthase gene from Toxoplasma gondii. J Biol Chem. 1993;268: 6269–6280.

22. Sharma H, Landau MJ, Vargo MA, Spasov KA, Anderson KS. First Three-Dimensional Structure of Toxoplasma gondii Thymidylate Synthase–Dihydrofolate Reductase: Insights for Catalysis, Interdomain Interactions, and Substrate Channeling. Biochemistry. 2013;52: 7305–7317. doi:10.1021/bi400576t

23. Sidik SM, Hackett CG, Tran F, Westwood NJ, Lourido S. Efficient genome engineering of Toxoplasma gondii using CRISPR/Cas9. PloS One. 2014;9: e100450. doi:10.1371/journal.pone.0100450

24. Shen B, Brown KM, Lee TD, Sibley LD. Efficient gene disruption in diverse strains of Toxoplasma gondii using CRISPR/CAS9. mBio. 2014;5: e01114–01114. doi:10.1128/mBio.01114-14

25. Ran FA, Hsu PD, Wright J, Agarwala V, Scott DA, Zhang F. Genome engineering using the CRISPR-Cas9 system. Nat Protoc. 2013;8: 2281–2308. doi:10.1038/nprot.2013.143

26. Winiger RR, Hehl AB. A streamlined CRISPR/Cas9 approach for fast genome editing in Toxoplasma gondii and Besnoitia besnoiti. J Biol Methods. 2020;7: e140. doi:10.14440/jbm.2020.343

27. Wang J, Tan Q, Chen J, Liu X, Di Z, Xiao Q, et al. Alkyl Hydroperoxide Reductase as a Determinant of Parasite Antiperoxide Response in Toxoplasma gondii. Oxid Med Cell Longev. 2021;2021: 1675652. doi:10.1155/2021/1675652

28. Pieperhoff MS, Pall GS, Jiménez-Ruiz E, Das S, Melatti C, Gow M, et al. Conditional U1 Gene Silencing in Toxoplasma gondii. PLOS ONE. 2015;10: e0130356. doi:10.1371/journal.pone.0130356

29. O’Shaughnessy WJ, Hu X, Beraki T, McDougal M, Reese ML. Loss of a conserved MAPK causes catastrophic failure in assembly of a specialized cilium-like structure in Toxoplasma gondii. Mol Biol Cell. 2020;31: 881–888. doi:10.1091/mbc.E19-11-0607

30. Briner AE, Donohoue PD, Gomaa AA, Selle K, Slorach EM, Nye CH, et al. Guide RNA Functional Modules Direct Cas9 Activity and Orthogonality. Mol Cell. 2014;56: 333–339. doi:10.1016/j.molcel.2014.09.019

31. Anders C, Niewoehner O, Duerst A, Jinek M. Structural basis of PAM-dependent target DNA recognition by the Cas9 endonuclease. Nature. 2014;513: 569–573. doi:10.1038/nature13579

32. Jinek M, Jiang F, Taylor DW, Sternberg SH, Kaya E, Ma E, et al. Structures of Cas9 Endonucleases Reveal RNA-Mediated Conformational Activation. Science. 2014;343: 1247997. doi:10.1126/science.1247997

33. Nishimasu H, Ran FA, Hsu PD, Konermann S, Shehata SI, Dohmae N, et al. Crystal structure of Cas9 in complex with guide RNA and target DNA. Cell. 2014;156: 935–949. doi:10.1016/j.cell.2014.02.001

34. Jiang F, Zhou K, Ma L, Gressel S, Doudna JA. STRUCTURAL BIOLOGY. A Cas9-guide RNA complex preorganized for target DNA recognition. Science. 2015;348: 1477–1481. doi:10.1126/science.aab1452

35. Jiang F, Taylor DW, Chen JS, Kornfeld JE, Zhou K, Thompson AJ, et al. Structures of a CRISPR-Cas9 R-loop complex primed for DNA cleavage. Science. 2016;351: 867–871. doi:10.1126/science.aad8282

36. Sundaresan R, Parameshwaran HP, Yogesha SD, Keilbarth MW, Rajan R. RNA-Independent DNA Cleavage Activities of Cas9 and Cas12a. Cell Rep. 2017;21: 3728–3739. doi:10.1016/j.celrep.2017.11.100

37. Zhang X-H, Tee LY, Wang X-G, Huang Q-S, Yang S-H. Off-target Effects in CRISPR/Cas9-mediated Genome Engineering. Mol Ther Nucleic Acids. 2015;4: e264. doi:10.1038/mtna.2015.37

38. Hu JH, Miller SM, Geurts MH, Tang W, Chen L, Sun N, et al. Evolved Cas9 variants with broad PAM compatibility and high DNA specificity. Nature. 2018;556: 57–63. doi:10.1038/nature26155

39. Nishimasu H, Shi X, Ishiguro S, Gao L, Hirano S, Okazaki S, et al. Engineered CRISPR-Cas9 nuclease with expanded targeting space. Science. 2018;361: 1259–1262. doi:10.1126/science.aas9129

40. Walton RT, Christie KA, Whittaker MN, Kleinstiver BP. Unconstrained genome targeting with near-PAMless engineered CRISPR-Cas9 variants. Science. 2020;368: 290–296. doi:10.1126/science.aba8853

41. Collias D, Leenay RT, Slotkowski RA, Zuo Z, Collins SP, McGirr BA, et al. A positive, growth-based PAM screen identifies noncanonical motifs recognized by the S. pyogenes Cas9. Sci Adv. 2020;6: eabb4054. doi:10.1126/sciadv.abb4054

42. Zhang Y, Ge X, Yang F, Zhang L, Zheng J, Tan X, et al. Comparison of non-canonical PAMs for CRISPR/Cas9-mediated DNA cleavage in human cells. Sci Rep. 2014;4: 5405. doi:10.1038/srep05405

43. Jiang W, Bikard D, Cox D, Zhang F, Marraffini LA. CRISPR-assisted editing of bacterial genomes. Nat Biotechnol. 2013;31: 233–239. doi:10.1038/nbt.2508

44. Jinek M, Chylinski K, Fonfara I, Hauer M, Doudna JA, Charpentier E. A programmable dual-RNA-guided DNA endonuclease in adaptive bacterial immunity. Science. 2012;337: 816–821. doi:10.1126/science.1225829

45. Wu X, Kriz AJ, Sharp PA. Target specificity of the CRISPR-Cas9 system. Quant Biol. 2014;2: 59–70. doi:10.1007/s40484-014-0030-x

46. Lin Y, Cradick TJ, Brown MT, Deshmukh H, Ranjan P, Sarode N, et al. CRISPR/Cas9 systems have off-target activity with insertions or deletions between target DNA and guide RNA sequences. Nucleic Acids Res. 2014;42: 7473–7485. doi:10.1093/nar/gku402

47. Duan J, Lu G, Xie Z, Lou M, Luo J, Guo L, et al. Genome-wide identification of CRISPR/Cas9 off-targets in human genome. Cell Res. 2014;24: 1009–1012. doi:10.1038/cr.2014.87

48. Alkan F, Wenzel A, Anthon C, Havgaard JH, Gorodkin J. CRISPR-Cas9 off-targeting assessment with nucleic acid duplex energy parameters. Genome Biol. 2018;19: 177. doi:10.1186/s13059-018-1534-x

49. Li B, Ren N, Yang L, Liu J, Huang Q. A qPCR method for genome editing efficiency determination and single-cell clone screening in human cells. Sci Rep. 2019;9: 18877. doi:10.1038/s41598-019-55463-6

50. Li R, Ba Y, Song Y, Cui J, Zhang X, Zhang D, et al. Rapid and sensitive screening and identification of CRISPR/Cas9 edited rice plants using quantitative real-time PCR coupled with high resolution melting analysis. Food Control. 2020;112: 107088. doi:10.1016/j.foodcont.2020.107088

51. Lee JS, Kallehauge TB, Pedersen LE, Kildegaard HF. Site-specific integration in CHO cells mediated by CRISPR/Cas9 and homology-directed DNA repair pathway. Sci Rep. 2015;5: 8572. doi:10.1038/srep08572

52. Mancini C, Messana E, Turco E, Brussino A, Brusco A. Gene-targeted embryonic stem cells: real-time PCR assay for estimation of the number of neomycin selection cassettes. Biol Proced Online. 2011;13: 10. doi:10.1186/1480-9222-13-10

53. Winzer P, Müller J, Aguado-Martínez A, Rahman M, Balmer V, Manser V, et al. In Vitro and In Vivo Effects of the Bumped Kinase Inhibitor 1294 in the Related Cyst-Forming Apicomplexans Toxoplasma gondii and Neospora caninum. Antimicrob Agents Chemother. 2015;59: 6361–6374. doi:10.1128/AAC.01236-15

54. Imhof D, Anghel N, Winzer P, Balmer V, Ramseier J, Hänggeli K, et al. In vitro activity, safety and in vivo efficacy of the novel bumped kinase inhibitor BKI-1748 in non-pregnant and pregnant mice experimentally infected with Neospora caninum tachyzoites and Toxoplasma gondii oocysts. Int J Parasitol Drugs Drug Resist. 2021;16: 90–101. doi:10.1016/j.ijpddr.2021.05.001

55. Winzer P, Anghel N, Imhof D, Balmer V, Ortega-Mora L-M, Ojo KK, et al. Neospora caninum: Structure and Fate of Multinucleated Complexes Induced by the Bumped Kinase Inhibitor BKI-1294. Pathog Basel Switz. 2020;9: E382. doi:10.3390/pathogens9050382

56. Longo MC, Berninger MS, Hartley JL. Use of uracil DNA glycosylase to control carry-over contamination in polymerase chain reactions. Gene. 1990;93: 125–128. doi:10.1016/0378-1119(90)90145-h

57. Costa J-M, Pautas C, Ernault P, Foulet F, Cordonnier C, Bretagne S. Real-Time PCR for Diagnosis and Follow-Up of Toxoplasma Reactivation after Allogeneic Stem Cell Transplantation Using Fluorescence Resonance Energy Transfer Hybridization Probes. J Clin Microbiol. 2000;38: 2929–2932. doi:10.1128/JCM.38.8.2929-2932.2000

58. Reischl U, Bretagne S, Krüger D, Ernault P, Costa J-M. Comparison of two DNA targets for the diagnosis of Toxoplasmosis by real-time PCR using fluorescence resonance energy transfer hybridization probes. BMC Infect Dis. 2003;3: 7. doi:10.1186/1471-2334-3-7

59. Donald RG, Roos DS. Insertional mutagenesis and marker rescue in a protozoan parasite: cloning of the uracil phosphoribosyltransferase locus from Toxoplasma gondii. Proc Natl Acad Sci. 1995;92: 5749–5753. doi:10.1073/pnas.92.12.5749

60. Nenarokova A, Záhonová K, Krasilnikova M, Gahura O, McCulloch R, Zíková A, et al. Causes and Effects of Loss of Classical Nonhomologous End Joining Pathway in Parasitic Eukaryotes. mBio. 2019;10: e01541–19. doi:10.1128/mBio.01541-19

61. Li Z, Zhang W, Chen Y, Guo W, Zhang J, Tang H, et al. Impaired DNA double-strand break repair contributes to the age-associated rise of genomic instability in humans. Cell Death Differ. 2016;23: 1765–1777. doi:10.1038/cdd.2016.65

62. Günther S, Matuschewski K, Müller S. Knockout Studies Reveal an Important Role of Plasmodium Lipoic Acid Protein Ligase A1 for Asexual Blood Stage Parasite Survival. PLoS ONE. 2009;4: e5510. doi:10.1371/journal.pone.0005510

63. Aravind L, Iyer LM, Wellems TE, Miller LH. Plasmodium biology: genomic gleanings. Cell. 2003;115: 771–785. doi:10.1016/s0092-8674(03)01023-7

64. Gardner MJ, Hall N, Fung E, White O, Berriman M, Hyman RW, et al. Genome sequence of the human malaria parasite Plasmodium falciparum. Nature. 2002;419: 498–511. doi:10.1038/nature01097

65. Smith RH, Chen Y-C, Seifuddin F, Hupalo D, Alba C, Reger R, et al. Genome-Wide Analysis of Off-Target CRISPR/Cas9 Activity in Single-Cell-Derived Human Hematopoietic Stem and Progenitor Cell Clones. Genes. 2020;11: E1501. doi:10.3390/genes11121501

66. Hoebeeck J, Speleman F, Vandesompele J. Real-time quantitative PCR as an alternative to Southern blot or fluorescence in situ hybridization for detection of gene copy number changes. Methods Mol Biol Clifton NJ. 2007;353: 205–226. doi:10.1385/1-59745-229-7:205

67. Stefano B, Patrizia B, Matteo C, Massimo G. Inverse PCR and Quantitative PCR as Alternative Methods to Southern Blotting Analysis to Assess Transgene Copy Number and Characterize the Integration Site in Transgenic Woody Plants. Biochem Genet. 2016;54: 291–305. doi:10.1007/s10528-016-9719-z

68. Tran PN, Tate CJ, Ridgway MC, Saliba KJ, Kirk K, Maier AG. Human dihydrofolate reductase influences the sensitivity of the malaria parasite Plasmodium falciparum to ketotifen – A cautionary tale in screening transgenic parasites. Int J Parasitol Drugs Drug Resist. 2016;6: 179–183. doi:10.1016/j.ijpddr.2016.09.003

69. Kim K, Boothroyd JC. Toxoplasma gondii: stable complementation of sag1 (p30) mutants using SAG1 transfection and fluorescence-activated cell sorting. Exp Parasitol. 1995;80: 46–53. doi:10.1006/expr.1995.1006

70. Soldati D, Boothroyd JC. Transient transfection and expression in the obligate intracellular parasite Toxoplasma gondii. Science. 1993;260: 349–352. doi:10.1126/science.8469986

71. Messina M, Niesman I, Mercier C, Sibley LD. Stable DNA transformation of Toxoplasma gondii using phleomycin selection. Gene. 1995;165: 213–217. doi:10.1016/0378-1119(95)00548-k

